# Silencing cuticular pigmentation genes enables RNA FISH in intact chemosensory appendagess

**DOI:** 10.1101/318535

**Authors:** Stefan Pentzold, Veit Grabe, Andrei Ogonkov, Lydia Schmidt, Wilhelm Boland, Antje Burse

## Abstract

Optical imaging of gene expression by RNA-fluorescent *in situ* hybridisation (FISH) in whole-mount sensory appendages of insects is often impeded by their highly pigmented cuticle. Since most chemical bleaching agents are incompatible with imaging fluorescent-labelled nucleotides, we developed a RNA interference-based method for clearing cuticular pigmentation that allows imaging of fluorescent mRNA in whole-mount appendages of insects. Silencing key genes of the tyrosine-derived pigmentation pathway by injecting dsRNA of *laccase2* or *tyrosine hydroxylase* in two leaf beetles species (*Chrysomela populi, Phaedon cochleariae*) resulted in clearance of the highly pigmented cuticle and in significant decreased light absorbance. Intact chemosensory appendages (palps, antennae and legs) from RNAi-cleared individuals were used to image expression and spatial distribution of antisense mRNA of two chemosensory genes (gustatory receptor, odorant-binding protein) via RNA FISH and confocal laser scanning microscopy. Imaging of these genes did neither work for RNAi-controls (ds*Gfp*) due to retained pigmentation, nor for FISH-controls using sense mRNA. Furthermore, we show that several chemical bleaching agents are not feasible with FISH, either due to significant degradation of polynucleotides, lack of clearing efficacy or long incubation times. Overall, silencing pigmentation genes is a significant improvement over bleaching agents allowing fluorescence imaging in whole-mount appendages and organs.

## Main text

Optical imaging in combination with *in situ* fluorescent labelling is a powerful method to elucidate spatiotemporal patterns of gene expression and to identify cellular circuits in biological systems^1^. For optimal image sharpness and resolution in fluorescence microscopy, samples should ideally be transparent. However, most tissues or appendages appear opaque and may be coloured by pigments. These obstacles prevent imaging at high depth – or any optical penetration at all – into tissue due to light absorption and scattering, respectively^2^. Thus, sharp imaging, especially deep into a tissue volume such as from whole-mount preparations, becomes strongly limited. Alternatively, physical serial sectioning in ultra-thin slices may be performed; however, correcting single slices for alignment, geometric distortion and staining variation is time-demanding and can be error-prone^3^. Non-sectioning approaches such as optical sectioning by confocal laser scanning (cLSM) or light sheet microscopy, largely preserve the innate 3-dimensional structure of biological tissues and organs, but require transparent samples^4^. Although a diverse set of novel tissue clearing techniques has been developed for some vertebrate species to achieve whole-organ or whole-body transparency enabling imaging of fluorescent labelled molecules deep into large tissue volume^4-12^, all current clearing methods come with specific limitations, e.g. they may affect structural and molecular integrity due to shrinkage or swelling of cells and tissue, quenching of fluorescence and/or altered antigenicity, among other issues^4,8^.

In many insect species, optical imaging is especially challenging, since pigmentation prevents fluorescence-based microscopy beneath the cuticle. Pigmentation and thus coloration can vary from colourless to yellowish or brownish to black depending on the amounts and types of melanin-like pigments incorporated^13^. Such pigments absorb light which results in decreased laser intensity and low optical quality as one penetrates deeper into tissue^3,14^. This bottleneck hampers deep imaging of intact tissue, appendages or whole bodies. Physical removal of the cuticle is less desirable due to inevitable destruction of surrounding tissue, especially in small specimen. Chemical clearing agents such as hydrogen peroxide (H_2_O_2_) bleach soft tissues such as brains, which allow reasonable fluorescent imaging of receptor neuron proteins by cLSM, but it results in tissue shrinkage such as in ants^15^ or spherically expanded ruptures in hawkmoths^14^. Other bleaching protocols for whole insects with brownish bodies require at least three weeks of incubation^16,17^. Since many insect species such as beetles often possess a particular thick, hard and pigmented cuticle, here we present a novel methodological strategy that combines both specific and effective clearing of cuticular pigmentation as well as preservation of molecular and cellular integrity. This approach allows optical imaging of fluorescence-labelled nucleotides in intact appendages or other whole-mount organs.

We take advantage of the proven functional importance of the enzymes tyrosine hydroxylase (TH) and laccase2 (Lac2, a phenol oxidase) in the beetle’s cuticular pigmentation pathway catalysing the first and following enzymatic steps^18,19^ (Fig. 1a). As expected and validated by qRT-PCR, *lac2* and *TH* were most expressed in the adult stage of the poplar leaf beetle (*Chrysomela populi*), especially in chemosensory appendages such as legs and antennae, which corresponds to the highest degree of pigmentation compared to other developmental stages and tissues, respectively (Fig. S1a). Silencing *TH* and *Lac2* by RNA interference (RNAi), i.e. injecting 80 or 140 ng double‐stranded RNA (dsRNA) of *Lac2* or *TH* per individual, resulted in significant clearing of cuticular pigmentation in comparison to RNAi-controls (injection of ds*Gfp*) in adult *C. populi* as well as the mustard leaf beetle *Phaedon cochleariae* (Fig. 1b). Injection of ds*Lac2* or ds*TH* in young pupae reduced pigmentation already in late pupae, but not in controls (Fig. S2). Hatching rate was similar among ds*Lac2* (68,5%, N=46), (ds*TH* 77,8%, N=22) and ds*Gfp*-control (65,8%, N=51) injected individuals; RNAi adults survived at least 8 days. Importantly, RNAi-based clearing was induced in chemosensory appendages such as antennae, legs and palps (Fig. 1b, c). Transcript levels were silenced significantly, i.e. by 94,5% and 98,2% for *CpopLac2* and *CpopTH* respectively in comparison to RNAi-controls, as validated by qRT-PCR (P=0.008, P=0.016; Fig. 1d).

**Figure 1.**
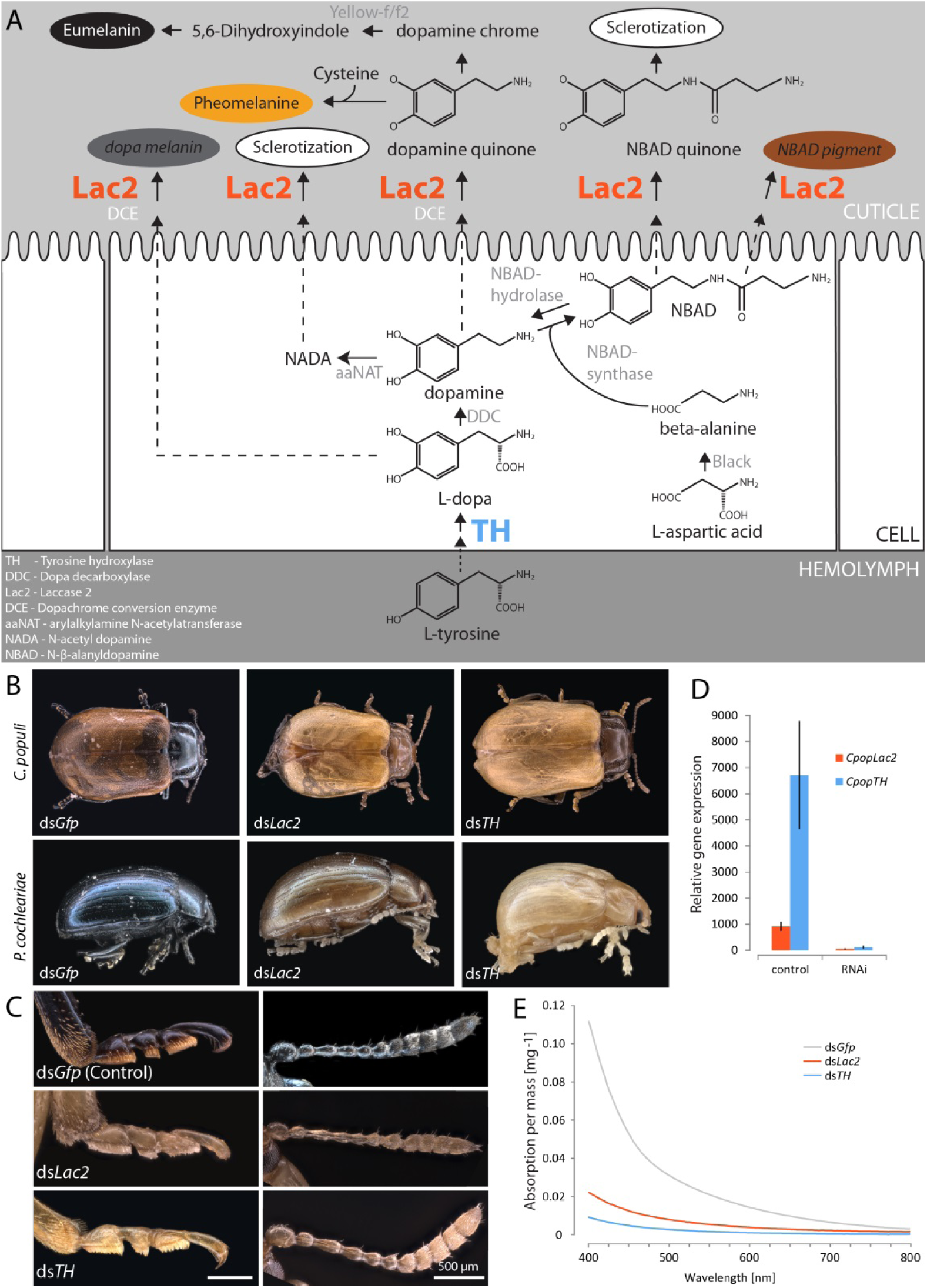
Silencing *TH* and *Lac2* genes by RNAi reduces cuticular pigmentation in adult leaf beetles. **A**. Tyrosine-derived cuticular pigmentation pathway in insects illustrating the involvement of tyrosine hydroxylase (TH, blue) and Laccase2 (Lac2, orange) enzymes with their respective substrates. The resulting pigments are coloured as proposed by^19,37,38^. **B:** Phenotypes of adult *C. populi* and *P. cochleariae* beetles silenced in *Lac2* and *TH* by RNAi (dsRNA injection) are less pigmented and thus lighter than RNAi-controls (ds*Gfp* injection). **C:** Detail on chemosensory appendages such as tarsi (left) and antennae (right) of RNAi and control individuals indicating cleared pigmentation. **D**: Efficacy of RNAi-mediated silencing of *CpopLac2* and *CpopTH* in comparison to RNAi-controls (injection of ds*Gfp*) as analysed by qRT-PCR. Differences in relative expression are statistically significant between the two treatments (*Lac2*: P=0,008; *TH*: P=0,016; Wilcoxon-Rank-Sum test) and refer to 94,5% and 98,2% respectively, transcript silencing in comparison to RNAi-controls. Bars represent ±s.e.m. **E:** Visible spectral changes measured by UV-Vis spectroscopy are associated with silencing *Lac2* or *TH* by RNAi in comparison to RNAi-controls (ds*Gfp*). Mean reduction in absorbance was 78% for ds*Lac2* and 91,4% for ds*TH*. Samples were macerated in acid and absorbance of the supernatant was measured from 400 – 800 nm.

To quantify clearing of cuticular pigmentation the visible spectrum of the supernatant of acid-macerated legs was measured by UV–Vis spectroscopy. The significant higher absorbance in the 400–475 nm region in RNAi-control samples, which is characteristic of dark-coloured products such as quinones from insect cuticles^13^, was missing in less-pigmented *lac2*- and *TH*-RNAi samples (Fig. 1e) showing a mean reduction in light absorbance of 78,0 and 91,4%, respectively. Therefore, we tested whether RNAi-cleared specimens are suitable for whole-mount RNA FISH and optical imaging by cLSM without chemical bleaching that would likely degrade mRNA.

As FISH targets, two distinct chemosensory gene families were chosen: a gustatory receptor (*CpopGR1*) and an odorant-binding protein (*CpopOBP13*). Both genes were most expressed in body appendages such the head (including palps), legs and antennae as shown by qRT-PCR (Fig. S1b). Similar to other gene expression studies in beetles^20-22^, *CpopGR1* had relative low overall expression levels, even in chemosensory organs (~10% of reference genes), whereas OBPs such as *CpopOBP13* are generally higher expressed as shown for the chemosensory appendages of *C. populi* (up to 15 times higher than reference genes). The RNAi treatment did not affect RNA FISH target sequences, i.e. the expression levels of *GR1* and *OBP13* did not differ between RNAi individuals (ds*Lac2* and ds*TH*) compared to RNAi-controls (P=0,39 for *CpopGR1*; P=0,73 for *CpopOBP13*, t-test). For RNA FISH, freshly dissected whole-mount chemosensory appendages (palps, antennae and legs) from RNAi-*Lac2* and RNAi-*TH* cleared beetles were incubated with biotin-labelled antisense mRNA (or sense mRNA as control) targeting either *CpopGR1* or *CpopOBP13* and combined with digoxigenin-labelled *CpopActin* as housekeeping gene. Importantly, gene expression of *CpopGR1* and *CpopOBP13* in intact appendages was successfully imaged via cLSM in those individuals that were silenced in *Lac2* or *TH* (Fig. 2a, Fig. S3a, b). For example, cells expressing *CpopGR1* were mainly proximal in the palp (Fig. 2a, i) and overall less abundant than those expressing *CpopOBP13* showing a broader distribution (Fig. 2a, ii). This is similar to the expression measured via qRT-PCR indicating relatively low or high expression for *CpopGR1* or *CpopOBP13*, respectively (Fig. S1b). Furthermore, in the antennae, *CpopOBP13* was expressed in the proximal and more distal parts (Fig. S3a), whereas in the tibia *CpopOBP13* was mainly expressed in proximal cells (Fig. S3b). Cuticular autofluorescence in RNAi-cleared organs remains to some extant after optical sectioning by cLSM (e.g. Fig. 2a i, ii), which is most likely caused by chitin^23^. In another study, H2O2-based quenching of autofluorescence was used to image microbial endosymbionts living in inner insect tissues such as the gut^16^; in general, many tissue clearing methods do not necessarily remove background autofluorescence^2^. In contrast to RNAi-cleared specimens, optical imaging of *CpopGR1* and *CpopOBP13* was not possible in RNAi-controls (ds*Gfp*) due to retained pigmentation that prevented optical penetration (Fig. 2a, iii). As further controls, using sense mRNA for FISH did not result in specific fluorescence for *GR1* or *OBP13* in any of the RNAi-treated beetles (ds*Lac2*, ds*TH* or ds*Gfp*-controls) (Fig. 2a, Fig. S3c). Nuclear DNA staining via fluorescent DAPI confirms cell integrity in RNAi-cleared tarsi of *C. populi* and *P. cochleariae* (Fig. 2b).

**Figure 2.**
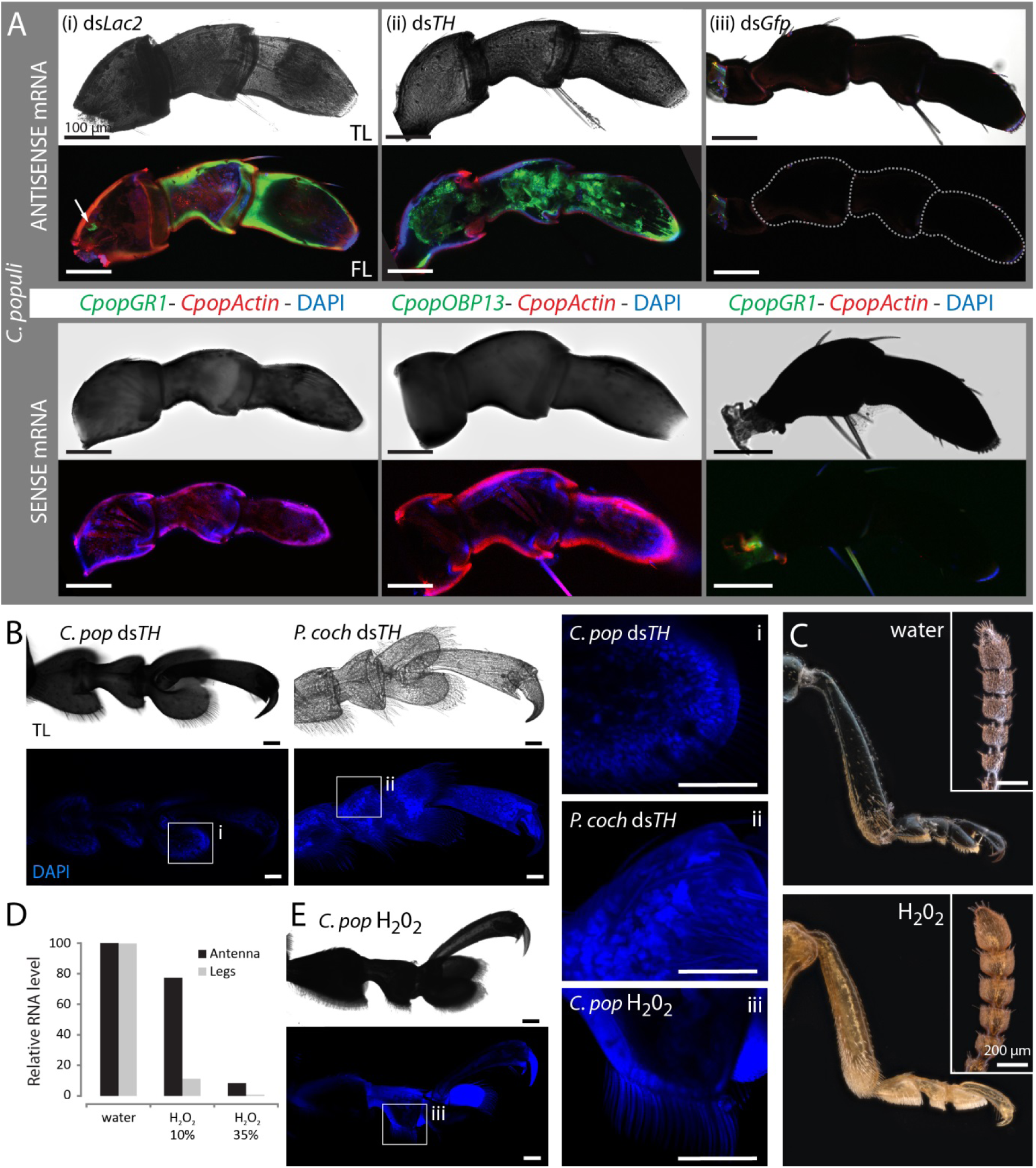
Specific clearing of cuticular pigmentation by RNAi enables RNA FISH in whole-mount appendages of leaf beetles. **A:** Maxillary palps from (i) *Lac2* or (ii) *TH*-silenced individuals with biotin-labelled mRNA of either a gustatory receptor (*CpopGR1*) or an odorant-binding protein (*CpopOBP13*) (both green) in combination with digoxigenin-labelled mRNA for *CpopActin* (red) as well as nuclear staining via DAPI (blue fluorescence). Due to remained pigmentation in RNAi-controls (ds*Gfp*, iii) RNA FISH using the same mRNA probes was not possible. Whereas antisense mRNA binds to the target sequence (upper panels), sense mRNA of *GR1* or *OBP13* as control does not bind (lower panels). TL - transmitted light microscopy (light panels), FL – fluorescence image from confocal laser scanning microscopy (dark panels). **B:** Tarsi of *C. populi* and *P. cochleariae* cleared by silencing *TH* and stained with DAPI targeting nuclear dsDNA confirms cellular integrity of RNAi treatments (i, ii). **C:** Incubation of antennae and legs of *C. populi* in H_2_O_2_ (35%) for 3 days clears cuticular pigmentation in comparison to water-controls, but **D**: results in significant degradation of total RNA in both organs. **E**: Tarsi incubated in H_2_O_2_ for 3 days could not be stained by DAPI indicating cellular damage by H_2_O_2_. Fluorescence is cuticular autofluorescence (iii).

To corroborate that RNAi-based clearing of cuticular pigmentation is the means of choice for using whole-mount organs in RNA FISH, we show that several bleaching chemicals are not feasible FISH, either due to significant degradation of polynucleotides, lack of clearing efficacy or long incubation times. We first tested H_2_O_2_ of which hydroxyl radicals and OOH-groups are known to destroy the cuticle-pigment melanin^24^. Whereas incubation of dissected body appendages in 35% H_2_O_2_ for 3 days reduced cuticular pigmentation in legs (Fig. 2c), it degraded 99,2% of the total RNA as measured by qRT-PCR using two house-keeping genes in comparison to water-incubated controls (Fig. 2d); similarly, antennae were chemically bleached resulting in degradation of 91,6% total RNA (Fig. 2c, d). Incubation of antennae and legs for 3 days in water (Fig. 2c) or 10% H_2_O_2_ did not reduce cuticular pigmentation, but degraded RNA (Fig. 2d). When incubating in alcoholic 6% H_2_O_2_, it took at least three weeks until bleaching of intact or bipartite antennae and legs (Fig. S4a). This is similarly long to bleaching brown-coloured, but relatively small insects such as lice, bat flies or aphids^16^. However, almost two third of total RNA (63,7%) was degraded despite prior to tissue fixation. When whole beetles were incubated in 35% H_2_O_2_ for five days, cuticular decolouration was evident, but partly incomplete, especially on the legs and wings that were also damaged (Fig. S4b). Consequently, cell integrity in legs and antennae was not retained in H_2_O_2_-bleached samples as indicated by unsuccessful DAPI staining (Fig. 2e). The cell-damaging characteristics of H_2_O_2_ resulting in degradation of total RNA and decreased fluorescence emission from green fluorescent proteins^25^, makes this bleaching method unsuitable for (RNA) FISH. This is especially important when targeting lowly expressed genes such as most insect GRs^20^. Other clearing agents such as methyl salicylate^3^ did clear soft tissues such as the brain of *C. populi*, after dehydration in an ethanol series, but not cuticular pigmentation (Fig. S5a). Similarly, a mixture of benzyl alcohol/benzyl benzoate did not clear cuticular pigmentation in antennae or legs (Fig. S5b).

RNAi-based clearing of cuticular pigmentation is a significant improvement compared to chemical bleaching agents, since it preserves molecular and cellular integrity and enables optical imaging of fluorescent-labelled nucleotides deep into tissue of intact whole-mount samples. This makes serial sectioning of organs redundant. We expect that our method also works with other, non-chemosensory genes regardless of whether they encode soluble proteins or membrane receptors, or whether they are lowly or highly expressed. Moreover, since RNAi-based suppression of *Lac2* or *TH* decreases cuticular pigmentation in different coleopteran^18,26-28^ and several other non-coleopteran insect species^29-34^, our method can be adapted to many other hexapod species enabling investigations beyond transparent embryos^35^ or ovaries^36^. Imaging beneath the RNAi-cleared cuticle may further benefit from light sheet microscopy that minimizes fluorophore bleaching and phototoxic effects. Finally, RNAi-based cuticle clearing complements existing toolkits such as by enabling *in vivo* calcium imaging, in contrast to chemical bleaching which does not maintain most life functions.

## Methods

### Specimen, RNA extraction and cDNA synthesis

Poplar leaf beetles (*C. populi*, L.,) were collected close to Dornburg (+51°00′52″, +11°38′17″) in Thuringia, Germany, where beetles were feeding on *Populus maximowiczii* x *Populus nigra* and reared in a climate chamber provided with fresh *P. nigra* twigs under 16:8 h L:D cycles at 20°C. Mustard leaf beetles (*P. cochleariae*, F.) were lab-reared on *Brassica oleracea* under same conditions. Both beetle species belong to the tribe Chrysomelini within the family Chrysomelidae. After homogenizing adult beetles using mortar and pestle in liquid nitrogen. RNA was purified using RNAqueous™ Total RNA Isolation Kit (Ambion) including DNase treatment following the manufacturer’s instruction. RNA concentration was measured on a NanoDrop™ One (Thermo Scientific). Synthesis of cDNA was carried out using SuperScript III Reverse Transcriptase and oligo(dT)20 (Invitrogen) following manufacturer’s protocol.

### Identification of candidate genes and molecular cloning

Orthologues of *Lac2* and *TH* in *C. populi* and *P. cochleariae* were identified by performing a BLAST search against the respective transcriptome^39,40^ using nucleotide sequences of *Lac2* (GenBank accession number: AY884061.2) and *TH* (EF592178) from *Tribolium castaneum* as template^18,28^. As targets for RNA FISH, a gustatory receptor (*CpopGR1*) and an odorant-binding protein (*CpopOBP13*) were identified from the transcriptome of *C. populi*^39^ by BLAST search using the template sequences CbowGR1 (GenBank: KT381521) and CbowOBP13 (KT381495) from the cabbage beetle *Colaphellus bowringi*^22^ (Chrysomelini, Chrysomelidae). The sequence of the house-keeping gene actin from *C. populi* was obtained from GenBank (JX122919.1). Gene specific primers were designed using Primer3 software (see table S1); a T7 promoter sequence (5’-TAATACGACTCACTATAGGGAGA-3’) was appended to forward and reverse primers of *Lac2* (*CpopLac2*, *PcoLac2*) and *TH* (*CpopTH*, *PcoTH*) to allow *in vitro* synthesis of dsRNA for RNA interference (see below). Therefore, PCR amplification was carried out using a proof-reading DNA polymerase (Phusion High-Fidelity, ThermoFisher). Amplicons were analysed via gel electrophoresis, purified using QIAquick PCR Purification (Qiagen) and sequenced. After RACE-PCR to verify full-length sequence using SMARTer RACE cDNA Amplification Kit (Clontech), *CpopGR1*, *CpopOBP13* and *CpopActin* were PCR-amplified without modification, sequenced and ligated into pCR-BluntII-TOPO (Invitrogen) posseesing opposing T7 and SP6 promoter sequences to allow *in vitro* synthesis from one template for generation of sense and antisense probes for RNA FISH (see below: Synthesis of labelled RNA FISH probes).

### RNA interference

#### Synthesis of dsRNA

Synthesis of dsRNA was carried out using MEGAscript^®^ RNAi Kit (Ambion) according to the manufacturer’s recommendations. In brief, PCR product with opposing T7 promoters flanking the target sequences (*Lac2*, *TH* from *C. populi* or *P. cochleariae*) was transcribed *in vitro* by incubating for 6 h at 37°C with T7-Polymerase, ATP, CTP, GTP and UTP including RNase inhibitor protein. Formation of dsRNA was induced by incubating the reaction mixture for 5 min at 75°C and then cooling to room temperature (RT). After nuclease digestion of single-stranded RNA and DNA, dsRNA was purified using filter cartridges and centrifugation. Finally, dsRNA was ethanol precipitated, resuspended in 0.9% physiological NaCl and adjusted to 1 µg/µl. The concentration and integrity of the dsRNA were determined by spectrometry (NanoDrop One, Thermo Scientific) and gel electrophoresis. The following lengths of dsRNA were generated: *Cpop*ds*TH*: 535 bp, *Cpop*ds*Lac2*: 548 bp, *Pco*ds*Lac2*: 521 bp, *Pco*ds*TH*: 544 bp (all excluding 23bp T7-promoter sequence, see table S1). DsRNA of *Gfp* (719 bp) was used as control.

#### Microinjection

Microinjections were performed using pulled borosilicate glass capillaries and Nanoliter 2000 injector (World precision instruments, Inc.) Injection was parasagittally between the pro- and mesothorax of either last larval instars or 3-day old pupae of *C. populi* and *P. cochleariae*. 12 biological replicates were run, i.e. a total of 48. Insects were briefly anesthetized on ice before and during injection. Each specimen was injected 20, 70 or 150 ng, respectively of total dsRNA, which corresponds to two times 9.2, 32.2 or 69.0 nl of *Lac2*, *TH*, *lac2*+*TH* or *Gfp*. Five to ten days after eclosure and rearing adult beetles on *P. nigra*, chemosensory appendages such as legs, antennae and palps were dissected for use in RNA FISH (see below). Additionally, efficacy of RNAi-mediated knockdown of *CpopLac2* and *CpopTH* was assessed using quantitative real-time PCR on cDNA derived from legs. Therefore, transcript abundance of *Lac2* or *TH* in ds*Gfp* control samples was set as baseline (100%) and compared to their expression in RNAi beetles. For statistical analysis, mean ct values of *Lac2* or *TH* from both groups (ds*Gfp* control versus RNAi) were compared for significant differences using Wilcoxon Rank-Sum test (SigmaPlot 12.0). Five biological replicates were analysed from both groups. For primers see table S1.

### Quantitative real time-PCR

Using qRT-PCR transcript abundance of (i) *Lac2* and *TH* was analysed in *C. populi* in different developmental stages such as larvae, pupae and adults, and in adult legs, antennae and wings (three to four biological replicates), (ii) *CpopGR1* and *CpopOBP13* in mentioned adult tissues such as gut, wings, thorax (excluding head and legs), legs, antennae and head including palps (six biological replicates). Data were quantified relative to the mRNA levels of the reference genes eukaryotic elongation factor 1-alpha (*EF1a*) and initiation factor 4A (eIF4A) using the 2− ^ΔΔCt^ –method^41^. Data were acquired on a CFX-96 Touch™ Real-Time PCR Detection System (Bio-Rad) using Brilliant III Ultra-Fast SYBR^®^ Green QPCR Master Mix (Agilent Technologies) and cDNA as template. Distilled water as template served as negative control. Two technical replicates were analysed; those with a Ct difference of >0.5 were repeated. To exclude effects of RNAi (ds*Lac2*, ds*Gfp*) on the expression level of *CpopGR1* and *CpopOBP13* (as RNA FISH targets, see below), their expression was compared to ds*Lac2* or ds*Gfp*-injected individuals by qRT-PCR on cDNA derived from of legs (five biological replicates per group). Statistical difference of the ^ΔCt^-values between the two groups was compared using t-test (SigmaPlot 12.0). Standard curves in five different 10-fold dilutions using plasmids containing the target gene were used to calculate amplification efficiency. For each primer pair sigmoidal amplification curves were obtained with single melt curve peaks. No signals were detected for the negative controls. For primers used see table S1.

### Whole-mount double RNA FISH

#### Synthesis of labelled RNA FISH probes

Plasmids containing target sequences (*CpopGR1*, *CpopOBP13*, *CpopActin*) flanked by opposing T7 and SP6 promoter sequences were linearized using NotI or BamHI. Subsequent *in vitro* transcription of labelled antisense or sense RNA was carried out using SP6 or T7 polymerases in the presence of biotin (GR1, OBP13) or digoxigenin (DIG; actin) following manufacturer’s procedure (RNA Labelling mix 10x, Roche). After incubation for 3 h at 37°C, RNA was precipitated using ethanol, dissolved in RNase-free water and diluted in hybridisation buffer (50% deionized formamide, 2× SSC, 0.2 mg/mL sonicated herring sperm DNA, 200 µg/ml yeast tRNA, and 10% dextran sulfate, water bidest). The following lengths were generated: *CpopGR1*: 390 nt and *CpopOBP13*: 513 nt (both biotin-labelled) and *CpopActin* (DIG-labelled) about 450 nt from original 1131 nt after incubation in hydrolysis buffer (80 mM NaHCO_3_ and 120 mM Na_2_CO_3_) at 60°C for 14,5 min (formula see ^36^).

#### Fixation and FISH procedure

Intact and whole palps (labial, maxillary), antennae and legs were freshly dissected from ice-chilled adult *C. populi* beetles. To improve penetration of labelled FISH probes, antennae and legs were cross-sectioned into a distal and proximal part (at the 5^th^ flagellomere and between tibia and femur, respectively). Samples were fixed in 4% paraformaldehyde in 1M NaHCO_3_ at 4°C for 24 h. Organs were washed in 1xPBS containing 0,03% TritonX100 for 1 min at RT and incubated in 0.2M HCl containing 0,03% TritonX100 for 10 min. After washing in 1xPBS with 1% TritonX100 for 1 min at RT, prehybridization was carried out by incubating samples in hybridisation buffer for 4 h at 4°C and for additional 6 h at 55°C. Hybridisation of labelled RNA probes to endogenous transcripts was performed at 55°C for 3 days. The following combinations were used: antisense or sense mRNA of *GR1* or *OBP13* (biotin-labelled) in combination with antisense of actin (DIG-labelled). After washing in 0.1xSSC with 0,03% TritonX100 four times at 60°C by horizontal shaking at 350 rpm, samples were blocked in 1%-blocking solution (Roche) diluted in TBS and 0,03% TritonX100 for 6h at 4°C. Detection of DIG-labelled probes was achieved by using an anti-DIG-conjugated alkaline phosphatase that dephosphorylates non-fluorochromic HNPP added to the sample into HNP that precipitates to RNA in the presence of Fast Red TR (HNPP Fluorescent Detection Set, Roche). Detection of biotin-labelled probes was achieved by streptavidin-coupled-horse radish peroxidase and tyramide signal amplification TSA fluorescein detection kit (Perkin Elmer NEL701A001KT). In both cases, antibody incubation was carried out for 3 days at 4°C followed by substrate incubation for 6 h. Intermediate washing steps were carried out three times for 10 min in TBS-0,05% Tween. Counterstaining of cellular dsDNA was done using DAPI (4′,6-diamidino-2-phenylindole) nucleic acid stain (Molecular probes) in 30 mM PBS and incubated for 30 min at RT in dark and three times washing in PBS. Finally samples were transferred onto microscope slides covered in Vectashield antifade mounting medium (Vector Laboratories).

### Microscopy and image processing

Fluorescent images were acquired with a ZEISS LSM 880 Confocal Laser Scanning Microscope using a 20x/0.8 Plan-Apochromat or 40x/1.2 C-Apochromat W (all ZEISS, Germany), respectively. Excitation of the fluorophores was conducted through a 405 nm laser diode, a 488 nm Argon laser and a 543 nm Helium-Neon laser (ZEISS, Germany). The systems spectral Quasar detector was setup to detect the fluorophores at 415-490 nm for DAPI, 490-561 nm for biotin-labelled probes and 551-735 nm for DIG-labelled probes. The pinhole was adjusted to 1 airy unit for the crucial channel of the *in situ* probe. Reflected light brightfield images from the whole beetles as well as their appendages were obtained on an AXIO Zoom V.16 (ZEISS, Germany). Images were processed using the following imaging software: ZEISS ZEN, Helicon Focus, Fiji and Adobe Photoshop CS.

### Absorbance measurement

Legs and antennae were ground in 6M HCl and centrifuged to pellet cell and tissue debris. Supernatant was used to measure absorbance from 400 to 800 nm per milligram ground tissue on a UV/VIS spectrophotometer (Jasco V-550).

### Chemical bleaching

Freshly dissected and fixed antennae and legs from *C. populi* were incubated in H_2_O_2_ (Sigma-Aldrich 18304) at different concentrations (aqueous 35.5%, 10%) or (alcoholic 6%) versus water or ethanol controls at RT. In addition, whole beetles were incubated in 35.5% H_2_O_2_ for 5 days at RT. For incubation in methyl salicylate freshly dissected heads of *C. populi* were dehydrated in an ethanol series, transferred in glass vials containing 1:1 ethanol:methyl salicylate for incubation for 1 h and finally substituted by 100% methyl salicylate (Sigma Aldrich M6752) for incubation for one week at RT during gentle shaking (350 rpm). After dehydrating further samples in a methanol series, they were incubated in a 1:2 mixture of BABB (Benzyl alcohol/benzyl benzoate) and incubated for three weeks.

### RNA degradation

Degradation of cellular RNA was analysed from antennae and legs (incubated for 3 days in 35.5% H_2_O_2_ or 6% alcoholic H_2_O_2_ in comparison to control incubations (water or ethanol) via qRT-PCR after RNA extraction and cDNA synthesis from three biological replicates (5-10 antennae or legs mixed) as described above. Therefore, mean ^ΔΔCt^-values of two reference genes (*EF1a* and *IF4a*) were compared between the two test and control treatments taking the latter values as baseline (100%) of mRNA abundance.

### Data availability

The data that support the findings of this study are available from the corresponding author upon reasonable request. Sequences generated in this study are deposited in GenBank (www.ncbi.nlm.nih.gov/genbank) under the following accession numbers: *CpopLac2*: MH253687; *PcoLac2*: MH253688; *CpopTH*: MH253689; *PcoTH*: MH253690; *CpopGR1*: MH253691; *CpopOBP13*: MH253692.

### Author contributions

S.P., W.B. and A.B. conceived and designed the study. S.P. performed RNA extractions, cDNA synthesis, identification of candidate genes, molecular cloning, qRT-PCR, FISH probe synthesis, fixation and RNA degradation assay. S.P. and A.O. performed chemical bleachings. V.G. carried out microscopic analyses by cLSM, processed images and constructed illustrations. A.O. analysed RNAi efficacy, performed dsRNA microinjection and FISH procedure. A.O. and A.B. analysed survival of RNAi beetles. L.S. synthesised dsRNA, did microinjections and prepared buffer solutions. S.P., A.O., V.G. and A.B. analysed the data. S.P. wrote the manuscript, all authors revised it.

### Competing interests

The authors declare no competing interests.

## Acknowledgements

S.P. acknowledges project funding from the European Union’s Horizon 2020 research and innovation programme under the Marie Skłodowska-Curie grant agreement No 705151. All authors are grateful to Vera L. Hopfenmüller for running replicates of FISH and to Tobias Becker for setting up the absorbance measurement. Further financial support from the Max Planck Society is acknowledged.

**Supplementary Figure S1.**
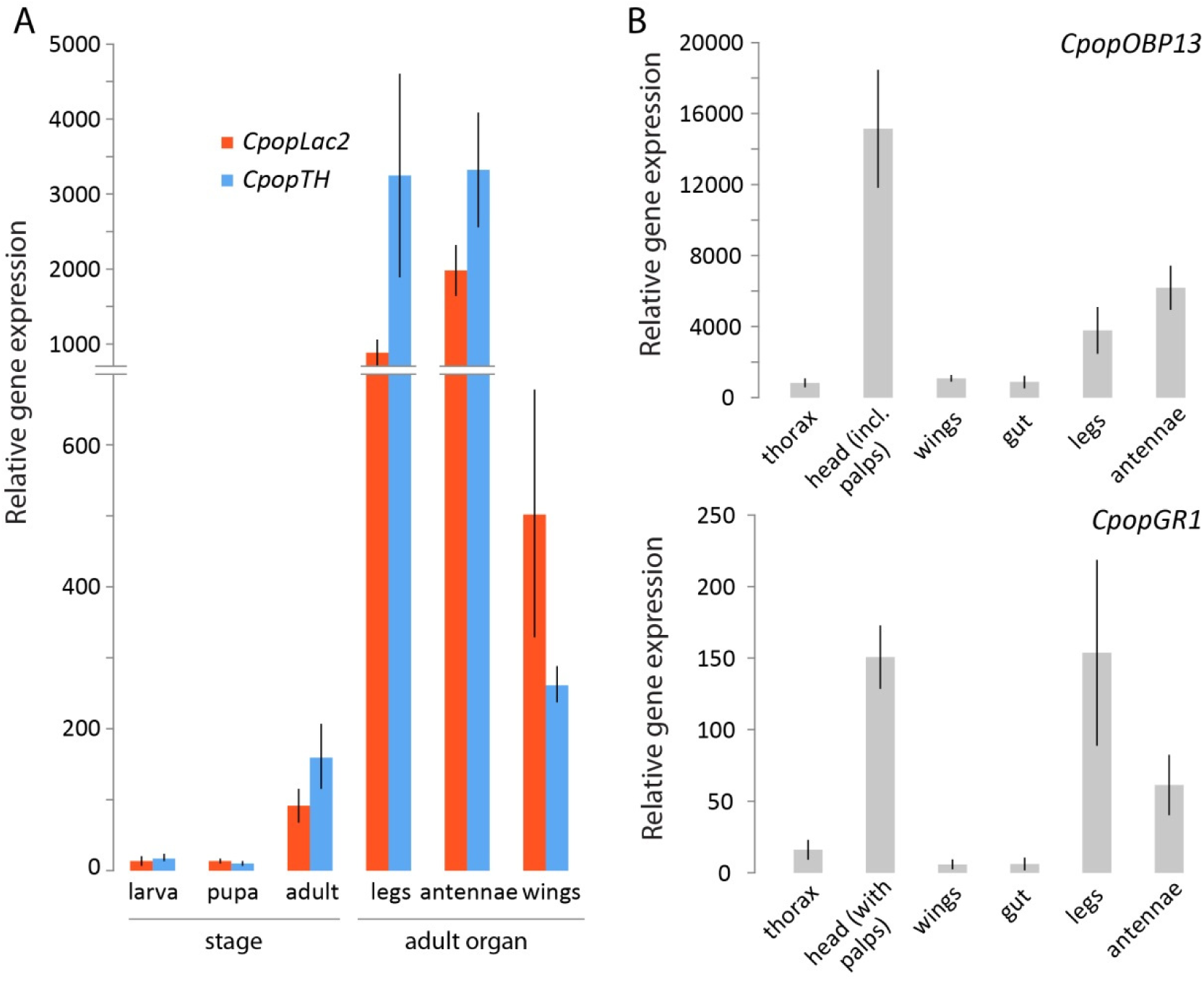
**A:** Expression of *Lac2* and *TH* as RNAi targets in untreated stages and adult tissues of *C. populi* as analysed by qRT-PCR. **B:** Expression of *CpopGR1* and *CpopOBP13* as RNA FISH targets in untreated tissues of adult *C. populi* as analysed by qRT-PCR. Bars represent ±s.e.m.

**Supplementary Figure S2.**
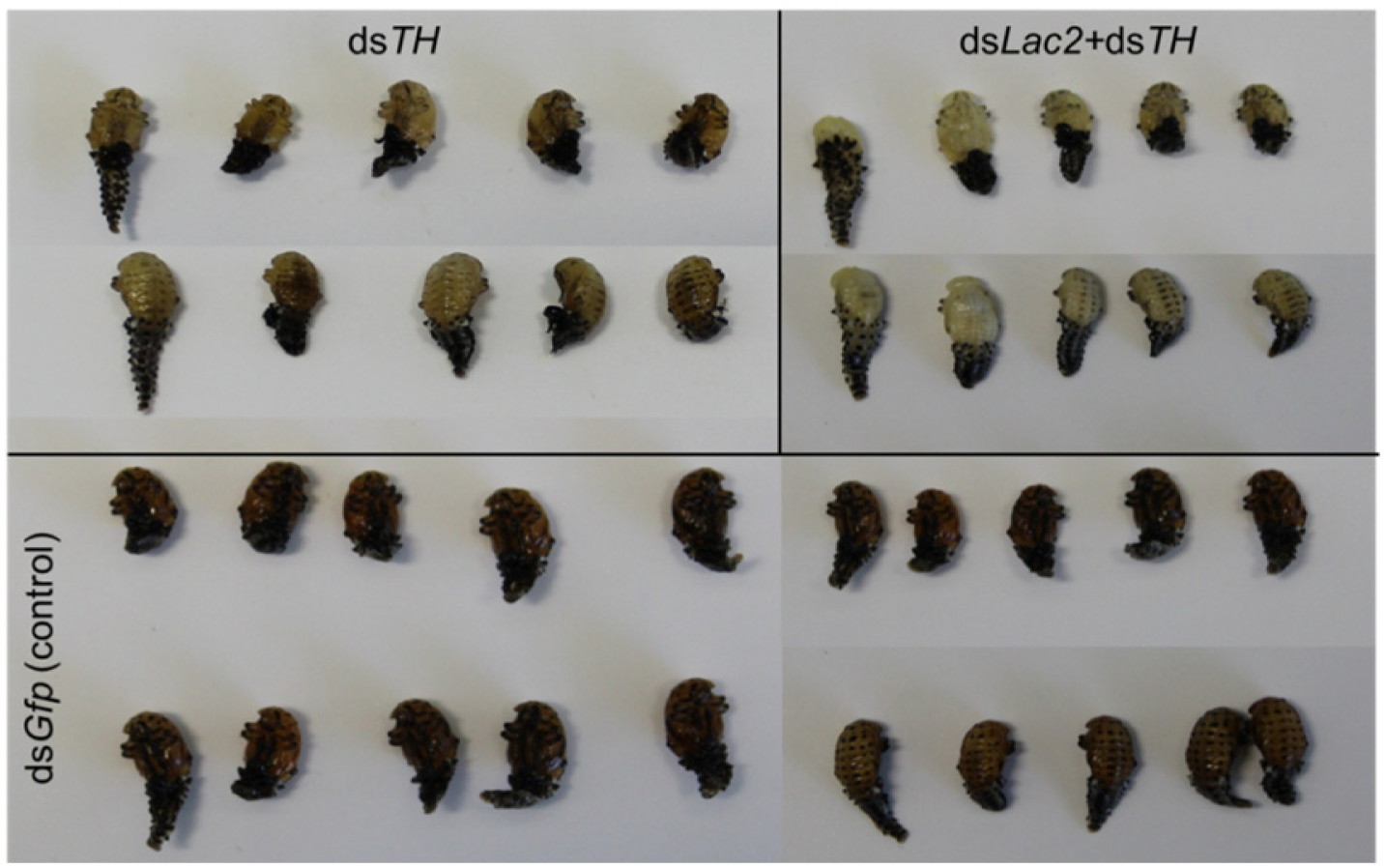
Young *C. populi* pupae of the same age were injected dsRNA targeting *TH* or *Lac2*+*TH* combined. RNAi-based clearing of cuticular pigmentation was evident in both cases in late pupae in comparison to darker RNAi-controls (ds*Gfp* injection).

**Supplementary Figure S3.**
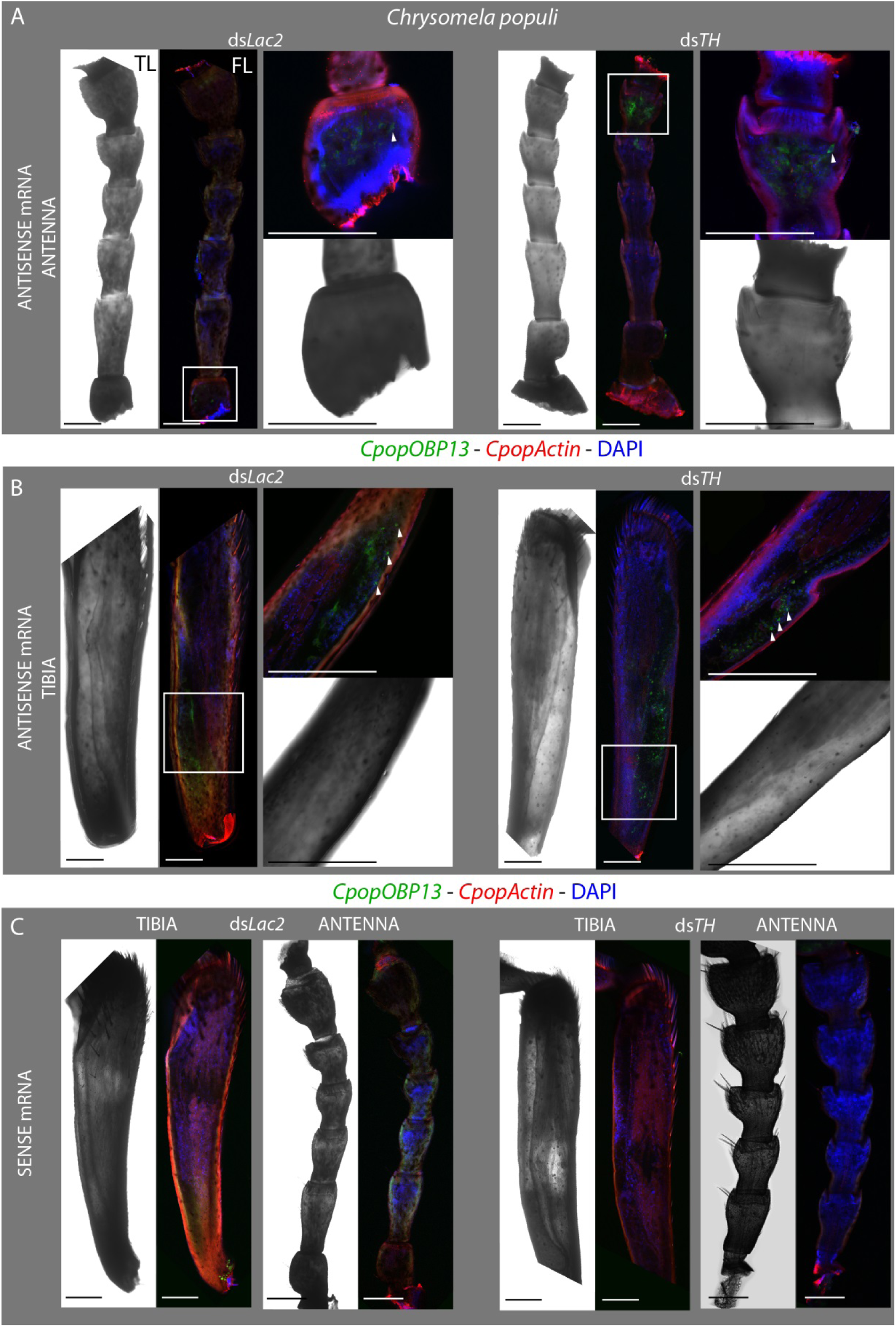
Clearing of cuticular pigmentation by RNAi enables FISH in whole-mount appendages of *C. populi*. **A:** Antennae and **B:** tibia from *Lac2* or *TH*-silenced individuals with biotin-labelled antisense mRNA of an odorant-binding protein (*CpopOBP13*) (green fluorescence) in combination with digoxigenin-labelled antisense mRNA for *CpopActin* (red) as well as DAPI staining nuclear dsDNA (blue). **C:** Using sense mRNA of *OBP13* as negative control in both RNAi-cleared appendages did not result in distinct green fluorescence, whereas positive controls (antisense *CpopActin*, DAPI) resulted in similar fluorescent signals as in A and B. Arrowheads indicate exemplary stained cells. Proximal parts of the organs are at the bottom, and distal part at the top of the image. TL - transmitted light microscopy (light panels), FL - fluorescence image from confocal laser scanning microscopy. Scale bar represents 200 µm.

**Supplementary Figure S4.**
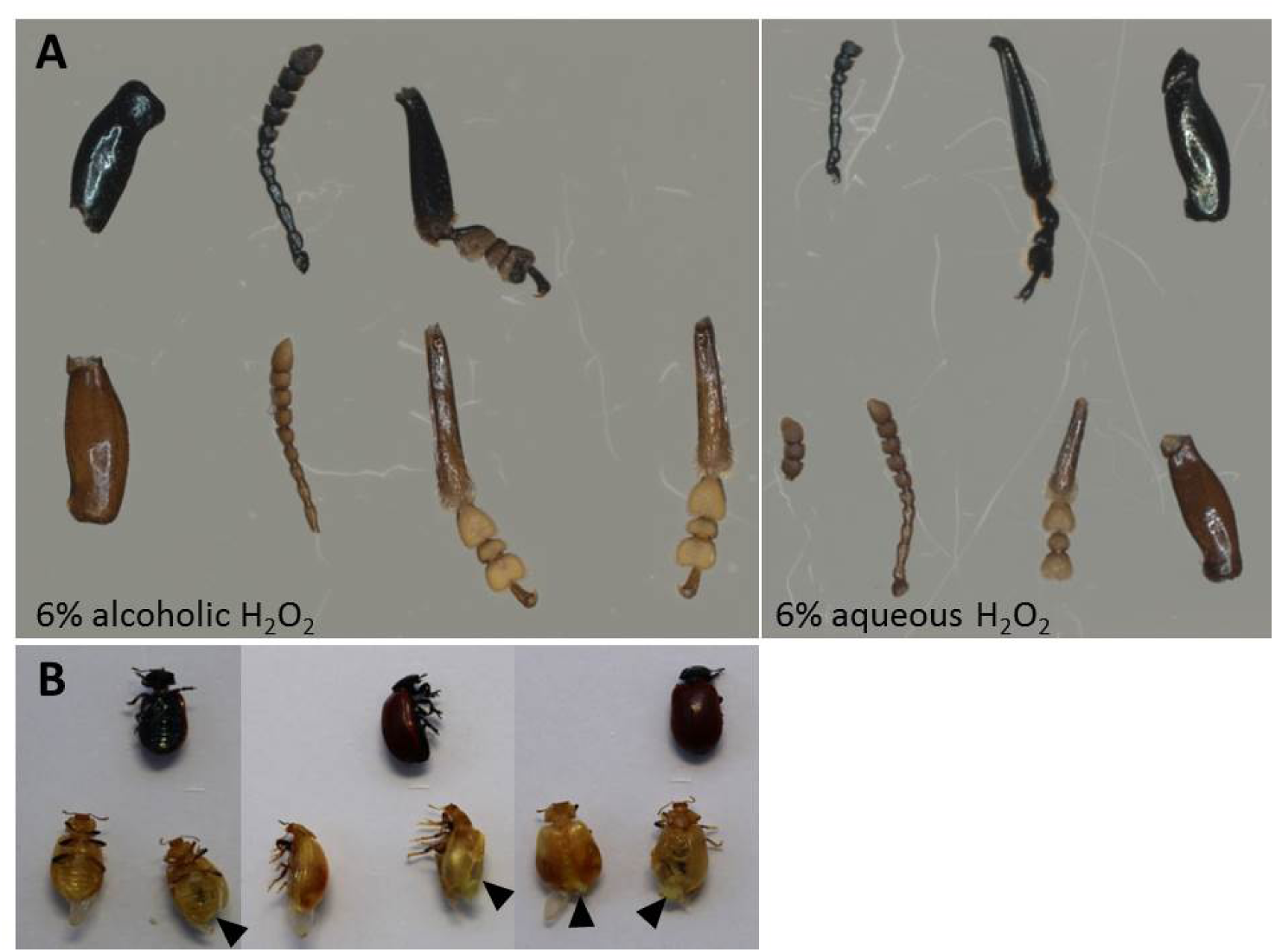
**A:** Incubation of antennae and bipartite legs from *C. populi* in 6% alcoholic or aqueous H_2_O_2_ for three weeks clears pigmentation in comparison to control incubations (ethanol or water), but degrades almost two third of total RNA (see text). **B:** Incubation of whole beetles in 35% H_2_O_2_ (bottom row) for five days results in cuticular decolouration (in comparison to controls, top row), which is however partly incomplete, especially on the legs while in some cases organs such as wings were damaged (arrowheads).

**Supplementary Figure S5.**
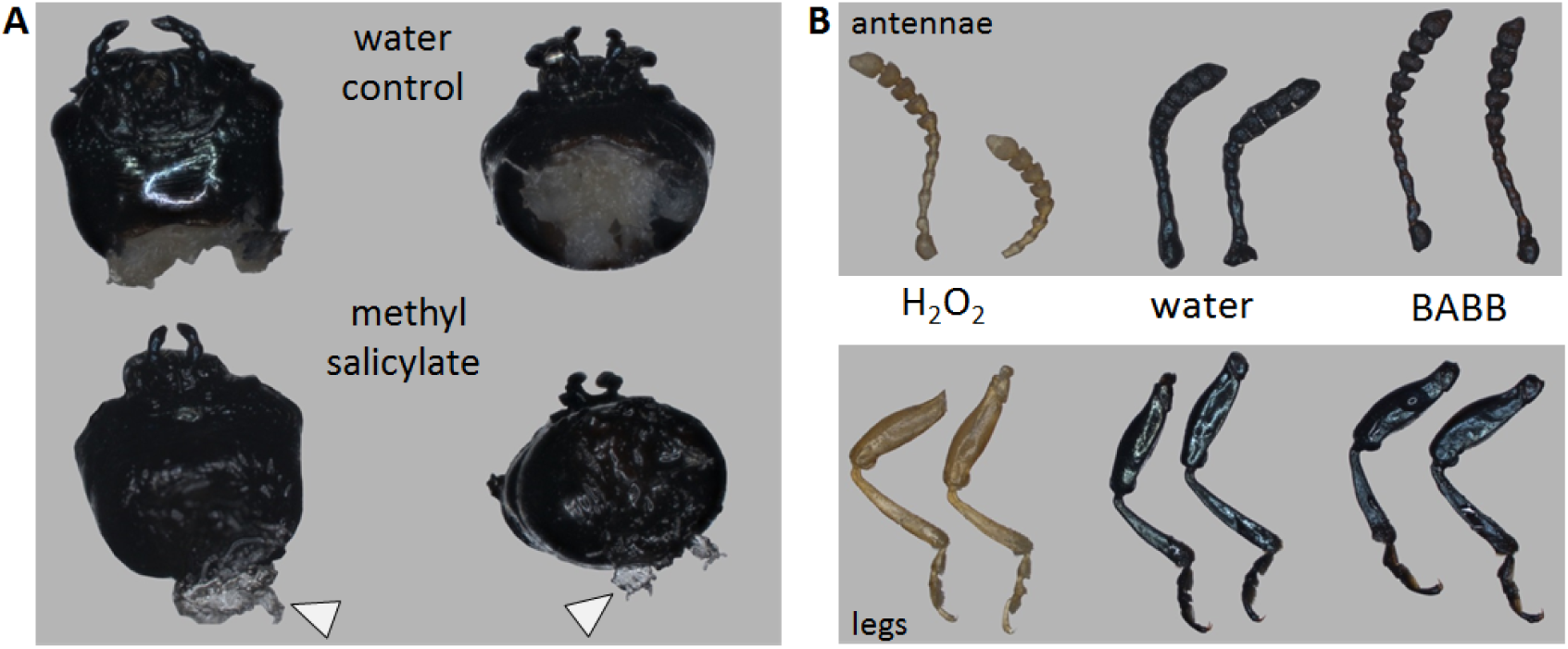
Incubation of chemosensory appendages of *C. populi* in **A:** methyl salicylate for one week cleared soft tissue such as the brain (arrows), but not the cuticle; **B:** BAAB (1:2 benzyl alcohol/benzyl benzoate) for three weeks did not clear cuticular pigmentation of antennae or legs in contrast to incubation in 35% H_2_O_2_. All incubations were done after fixating and dehydrating samples.

**Supplementary Table S1:**
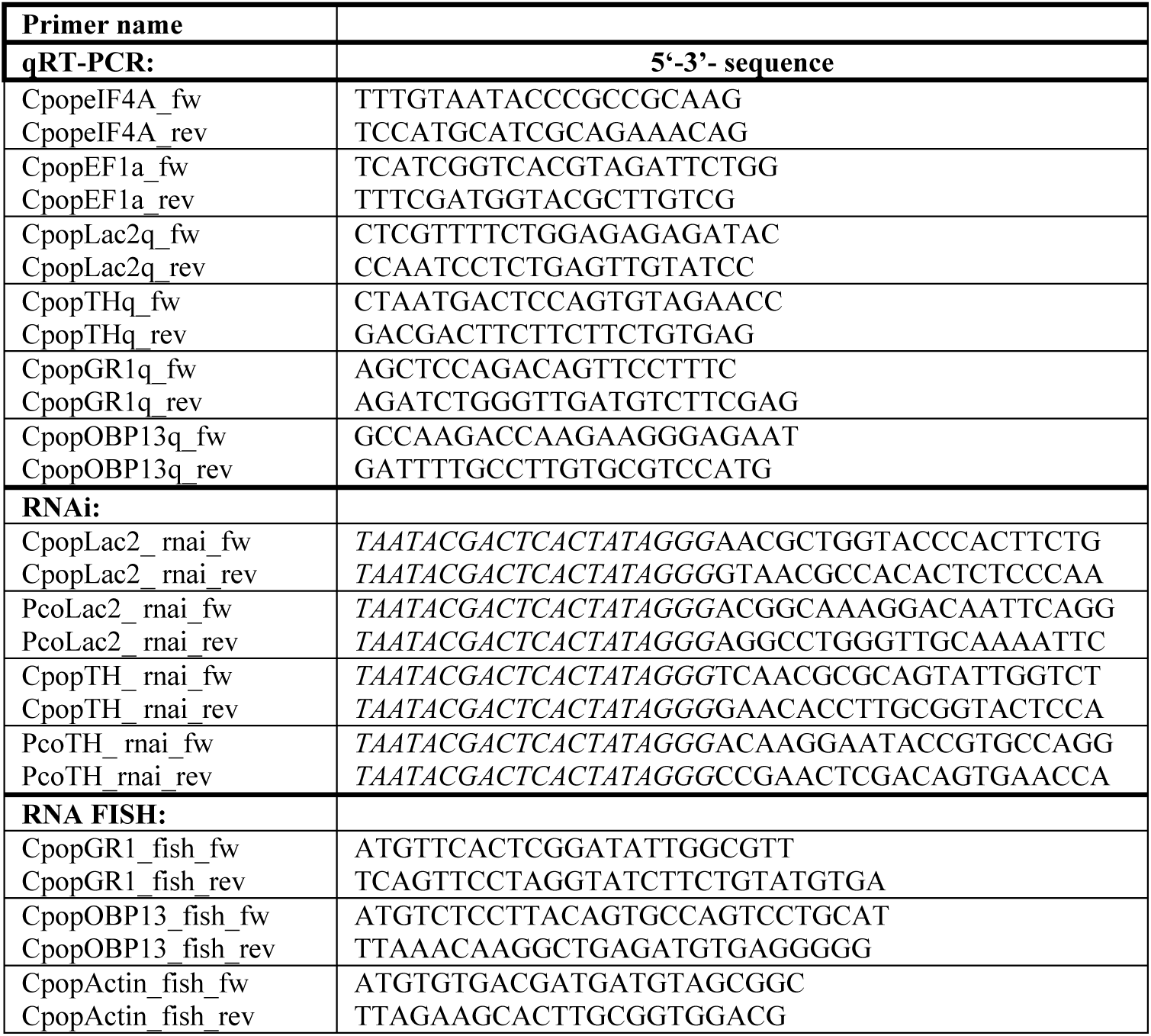
Primer sequences used for qRT-PCR, RNAi and RNA FISH. Cpop – *Chrysomela populi*; Pco – *Phaedon cochleariae*. T7 promoter sequence in italics. eIF4A – eukaryotic initiation factor 4 alpha; EF1 – eukaryotic elongation factor 1; Lac2 – laccase2; TH – tyrosine hydroxylase; GR – gustatory receptor; OBP – odorant binding protein; fw – forward; rev – reverse.

